# Convergence of Cortical and Thalamic Origins of Free Behavior Modulation of Mouse Primary Visual Cortex

**DOI:** 10.64898/2026.01.07.698022

**Authors:** Peijia Yu, Ha Yun Anna Yoon, Yuhan Yang, Yunlong Xu, Olivia Gozel, Gengshuo John Tian, Na Ji, Brent Doiron

## Abstract

The quality of sensory information flow in the brain may be affected by uninstructed free animal behavior. However, the specific circuit pathways that merge sensory and free behavior remain obscure. In mouse primary visual cortex (V1), we combine two-photon calcium imaging of cortical neurons and thalamo-cortical LGN (lateral geniculate nucleus) boutons with simultaneous measurements of facial movements. When controlling for spurious time-structured ‘nonsense’ correlations, we observed representations of both eye movements and non-ocular facial movement features in both LGN bouton and V1 cortical neuron activity during grating input stimulation. Further, the correlation between V1 neurons and facial movement is larger for grating stimulation compared to blank stimulation, likely due to the integration of LGN inputs and modulation of V1 from higher brain centers during grating stimulation. Together, our results suggest a convergence of uninstructed non-visual signals from a persistent top-down pathway and a stimulus-gated bottom-up pathway in primary visual cortex.

## Introduction

A growing body of work has charted how various brain ‘states’ and behavior shape sensory processing, as observed across diverse spatial-temporal scales, cortical areas, and species (Urai et al., 2022). In rodent primary visual cortex (V1), neural activity exhibits widespread correlations with non-visual behavioral variables such as the level of arousal (Reimer et al., 2014; McGinley et al., 2015; Vinck et al., 2015; Shimaoka et al., 2018; Raut et al., 2025), locomotion (Niell and Stryker, 2010; Busse, 2018; Musall et al., 2019; Schneider, 2020; Salkoff et al., 2020), and notably, facial movements (Stringer et al., 2019; Dolensek et al., 2020; Syeda et al., 2024). A prevailing view links these correlations to top-down inputs from higher-order areas that carry information about internal brain states such as attention, expectation, and goals during task engagement (Harris and Mrsic-Flogel, 2013; McGinley et al., 2015; Keller and Mrsic-Flogel, 2018). In parallel, early subcortical stages are also state modulated, with recent studies implicating the retina (Liang et al., 2020; Schröder et al., 2020; Muller et al., 2023; Lapanja et al., 2025) and the dorsal lateral geniculate nucleus (dLGN) (Erisken et al., 2014; Roth et al., 2016; Aydın et al., 2018; Nestvogel and McCormick, 2022; Spacek et al., 2022; Socha et al., 2024). Yet the origins and underlying mechanisms of these early stage effects, and how they coordinate with top-down modulation in cortex, remain unresolved.

In our study we collect *in vivo* calcium imaging in thalamo-cortical axonal boutons and cortical neurons in awake mouse V1, together with simultaneous video recordings of facial movement. Our goal is to use facial movement to quantify free, uninstructed behavior and measure its influence on the activity of cortical and subcortical responses. An important, but often overlooked, statistical caveat in identifying neural-behavioral correlations is that sequential trials are not independent samples, and long timescale auto-correlated noise in each variable will result in spurious cross-correlations termed ‘nonsense correlations’ (Yule, 1926; Harris, 2020; Elber-Dorozko and Loewenstein, 2018). Applying appropriate controls to discount nonsense correlations, we nevertheless find a significant behavioral correlate in the activity of feedforward thalamo-cortical dLGN boutons, but only during visually stimulated periods. Through comparative analysis that separates low-dimensional features of facial movement, this feedforward correlation partly, but not fully, attributes to eye motion. We show that the correlation between free behavior and cortical neuron activity is, unlike bouton activity, present even during blank stimulus periods, likely due to top-down pathways carrying non-visual inputs to V1. This correlation is increased during visual stimulation, consistent with dLGN providing a second, bottom-up pathway for behavioral inputs gated by the presence of a visual input. Finally, the encoding of visual stimuli and facial movement in V1 population activity space is intertwined, while the identity of visual stimulus can be perfectly read out without inter-ference by avoiding the subspace that encodes facial movement. Taken together, these results support a convergence of feedforward bottom-up and feedback top-down sources of facial movement modulation in mouse V1.

## Results

### Data acquisition: simultaneous *in vivo* imaging in mouse V1 and facial movement recording

We used *in vivo* two-photon calcium imaging to investigate how facial movements are represented in mouse V1. In separate measurements we recorded population Δ*F/F*_0_ activity of cortical neurons and/or afferent thalamo-cortical boutons from dLGN in layer (L) 2/3 and L4 of V1 from awake, head-fixed mice (Figure 1A, 1B) (See Methods). The recording of neurons and boutons in part of our L2/3 datasets (*n* = 5*/*7) were simultaneous (as in Figure 1A), while the neuronal and bouton recordings from L4 were collected separately. The visual stimulus consists of 1000 repeated cycles of a blank period and a stimulus period – the stimulus periods presented drifting grating stimuli with one of four different directions in random order (with 250 repeats for each direction); and the blank periods have gray stimulus with same average illuminance as the drifting grating stimuli.

**Figure 1:**
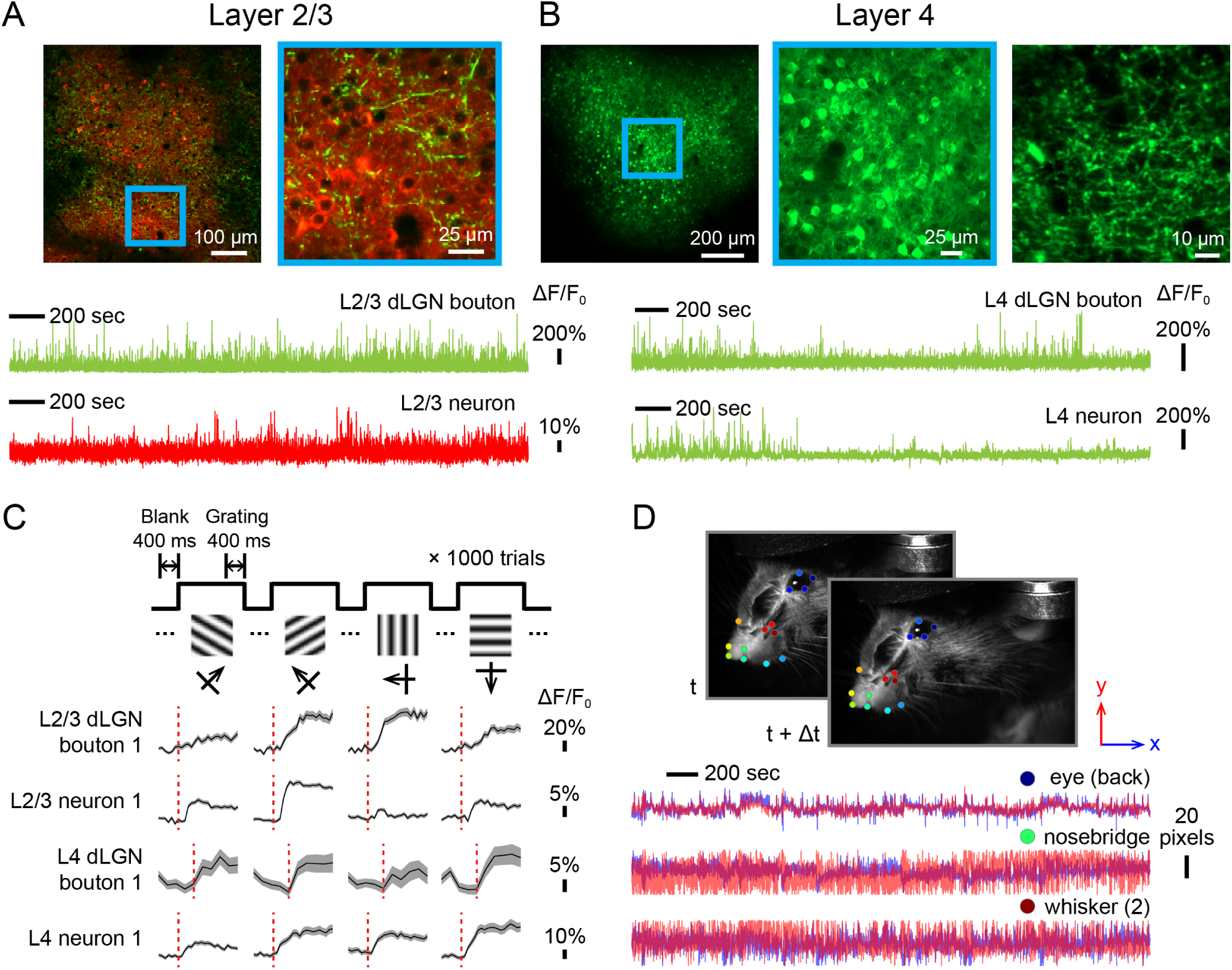
*In vivo* calcium imaging of population activity of cortical neuron or dLGN thalamo-cortical bouton in mouse V1 with simultaneous recording of facial movement. (A) Above: Example fields of view of *in vivo* calcium imaging of jRGECO1a+ neurons (red) and GCaMP6s+ thalamic boutons (green) in L2/3 of V1 in head-fixed awake mouse. Below: Δ*F/F*_0_(*t*) calcium traces of example boutons and neurons. (B) Above: Example fields of view in L4, with GCaMP6s+ neurons and thalamic boutons (green). (C) Above: Schematic of visual stimulus: repeated cycles of blank period and grating stimulus period, with drifting gratings of 4 different directions presented in random order. Trial responses were defined as the average response in non-overlapping 400 ms windows as indicated. Below: The Δ*F/F*_0_ transient traces over all frames in blank - grating cycle, averaged over all 250 repeated cycles for 4 different grating directions. Same example neurons and boutons as in (A) and (B). Red dashed lines indicate the transition frame from blank to grating period. Shadows represent standard error. (D) Above: Two example consecutive frames (indicated as *t* and *t*+Δ*t*) from a video recording of a mouse’s facial movement, acquired simultaneously with calcium imaging. In total, 14 facial keypoints (marked with distinct colors) are tracked by a fine tuned Facemap model (Syeda et al., 2024). Below: traces of horizontal (*x*, blue) and vertical (*y*, red) locations of three example keypoints.

Δ*F/F*_0_ responses in non-overlapping 400 ms windows were averaged as trial-responses for blank period and for grating periods in each direction (Figure 1C). Simultaneous with the neuronal/bouton recordings, the facial movements of mice were video recorded and tracked with a fine tuned FaceMap model (Syeda et al., 2024), yielding trajectories of x and y locations of 14 keypoints on the face (e.g. eye boundary, nose, whiskers). This provided a 28-dimensional representation of facial movement (Figure 1D). Given these datasets, we aimed to quantify the relationship between the trial-to-trial variability in facial movement and population activity of neuron or bouton with a linear regression model (see Methods). Because these datasets were acquired with identical stimulus structure, we could directly compare the correlation of facial movement in cortical activity and thalamo-cortical input. The facial movement was not associated with any cognitive tasks or rewards, and thus reflects the uninstructed, free behavior of the animal.

### Assessing ‘nonsense-free’ facial movement correlations

Linear regression is widely used to measure the correlation between neural activity and various types of behavior variables. Standard linear regression assumes all trials to be independent samples. However, when regression is completed on time series – which is common in neurophysiological and behavioral signals – it is vulnerable to so-called *nonsense correlations*. The idea of nonsense correlations was first introduced nearly a century ago (Yule, 1926) to describe the artifactual relationship between mortality rates and the fraction of marriages conducted by the Church of England, and more recently to illustrate comical spurious correlations between cryptocurrency prices and Neuropixel recordings in mice (Meijer, 2021), and between stock market value and rat neuronal activity (Marzullo et al., 2016). In brief, if two signals both contain significant trends over time (i.e auto-correlated), they can appear to be cross-correlated even without a bona fide genuine correlation. Unless further evidence shows that these trends over time should be modeled as signals rather than noise, this apparent ‘nonsense’ correlation can lead to an overestimate of actual correlations between neuronal activity and behavior. Fortunately, recent work has proposed several practical methods to detect nonsense correlations in neuronal data (Harris, 2020), with performance that depends on the temporal statistics of the data.

We first computed the auto-correlation structures in our neuronal and behavioral recordings. Across all separate datasets (neuron or bouton; in L2/3 or L4) the population Δ*F/F*_0_ activity show weak auto-correlation, while the auto-correlated memory in the dynamics of facial movement are significant (Figure 2A, left v.s. right). Furthermore, these auto-correlations asymptote to near zero in time lag, indicating robust stationary fluctuations rather than any long-term, systematic drifts. Importantly, for all datasets this decay occurs within the time lag of a single session (200 repeated cycles lasting 520 ∼ 790 sec.; where a full experiment contains five consecutive experimental sessions) (Figure 2A, red dashed lines), supporting a working assumption that the time dependence of both neural/bouton activity and facial movement are approximately statistically identical across different sessions, i.e. recorded under the same conditions.

**Figure 2:**
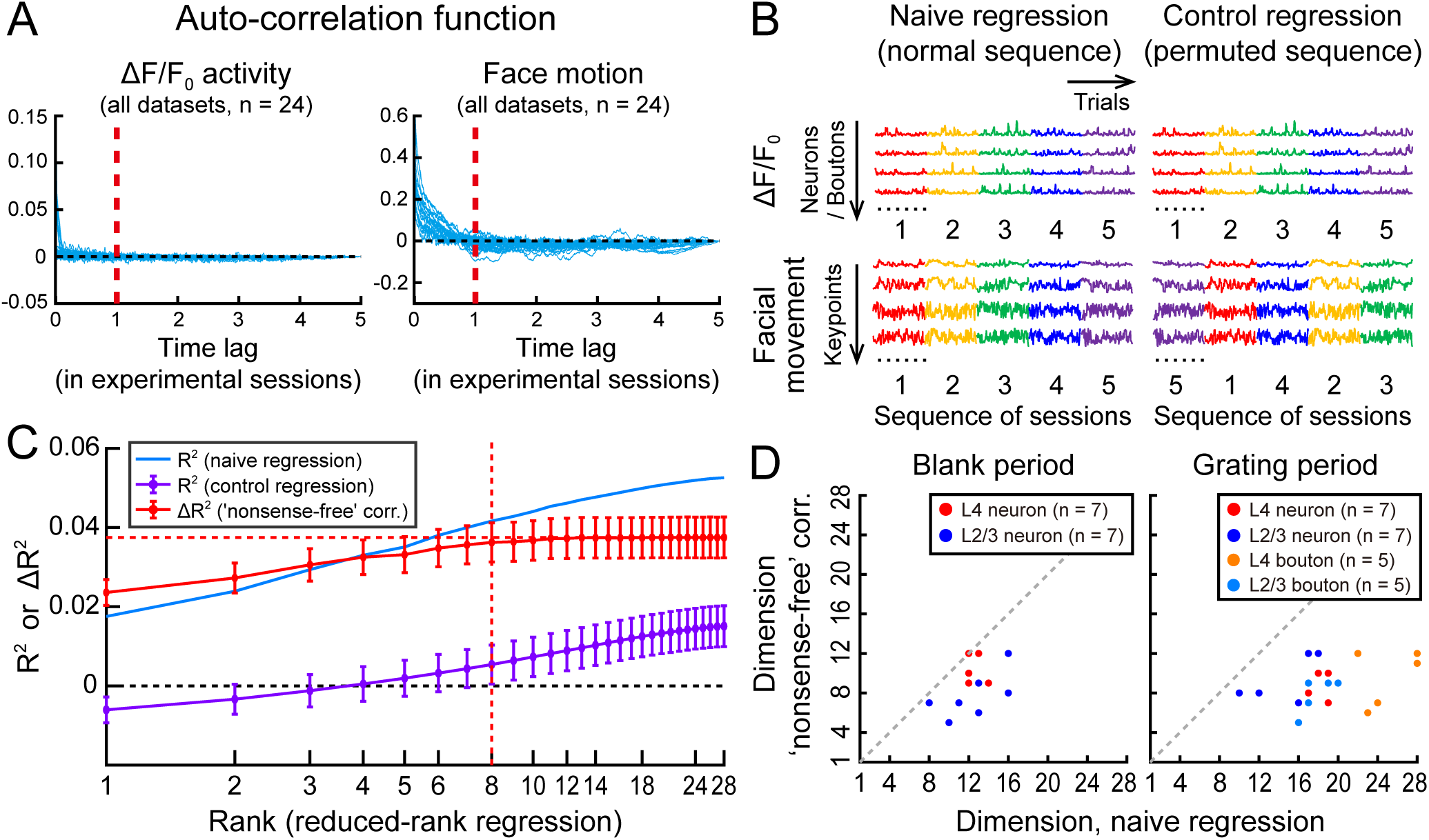
Attaining ‘nonsense-free’ correlation between Δ*F/F*_0_ activity and facial movement. (A) Auto-correlation functions of Δ*F/F*_0_ activity and facial movement as a function of time lag (recording frames), with all frames organized into 5 consecutive experimental sessions (each session lasts 200 repeated cycles lasting 520 ∼ 790 sec.). Each curve represents one dataset. For all datasets the auto-correlation asymptotes to near 0 within the duration of a single session (red dashed line), indicating approximately identical temporal statistics across sessions. (B) For data composed of multiple sessions recorded under identical conditions, the session permutation method provides an estimate of nonsense correlation. Left: a näıve regression contains both genuine and nonsense correlation. Right: in a control regression that permutes the sequence of sessions, genuine correlation is lost while nonsense correlation remains. Colors indicate sessions. (C) *R*^2^ from reduced-rank regression between Δ*F/F*_0_ activity and facial movement of one example dataset. The difference between näıve regression (blue) and permuted control regressions (purple), Δ*R*^2^, reflects the genuine, ‘nonsense-free’ correlation (red). Note that only ‘nonsense-free’ correlation saturates at a limited rank (horizontal and vertical red dashed lines). Error bars show standard deviation across all 5! − 1 = 119 permutations. (D) The effective dimensionality (define by the minimum rank that attains 95% of maximal correlation) in nonsense-free, genuine correlations are significantly lower than those in näıve regressions, for both blank and grating stimulus periods (Note that dimensionality of blank period is not applicable for boutons due to zero Δ*R*^2^, as in Figure 3A).

Given this stationary property, the session permutation control (Harris, 2020) was adopted to quantify the nonsense correlation in our datasets. A näıve regression between Δ*F/F*_0_ and facial movement contains both genuine and nonsense correlations (Figure 2B, left). For a control, we permuted the auto-correlated facial movement data in a special way: the order of trial within each session was kept intact but the sequence of five session blocks was permuted (Figure 2B, left v.s. right). This permutation breaks the genuine trial alignment while preserving the exact within-session temporal statistics that give rise to nonsense correlations. Thus, the gap between the näıve regression and the average of all possible permuted regressions (across all 5! − 1 = 119 non-identity orders), Δ*R*^2^, indicates the genuine behavioral correlation (Figure 2C).

Not only can nonsense correlations lead to over-estimation of correlation, they can also over-estimate the effective dimensionality (or rank) of the correlation between facial movement and neuron / bouton activity in reduced-rank regression. In an example dataset (during grating stimulus periods), genuine correlation asymptotes at around 8 dimensions (by attaining 95% of the maximal Δ*R*^2^), while the näıve regression does not asymptote and continues to rise up until the full rank (Figure 2C). For all datasets during blank or grating stimulus periods, genuine correlations show lower dimensionality than that of näıve regressions (Figure 2D). In sum, the temporal auto-correlation in our datasets lead to sizable nonsense correlations between neuronal activity and facial movements. However, when these spurious correlations are corrected, there remains significant genuine correlations with an effective dimension that is lower than the possible upper bound (28 dimensions).

### Facial movement correlation in feedforward visual pathway and V1 cortical neuron population

We summarize the ‘nonsense-free’ correlation, Δ*R*^2^, between facial movement and Δ*F/F*_0_ activity for thalamo-cortical bouton (Figure 3A) and cortical neuron (Figure 3B) activity in L2/3 and L4, during blank stimulus periods versus that during grating stimulus periods (x and y axes in Figure 3A and 3B, respectively). For each dataset we report the saturated maximal value of Δ*R*^2^ in reduced rank regression (as in Figure 2C), and we include only neurons or boutons that are responsive to visual stimuli (see Methods). All reports during grating stimulus periods are averaged across 4 stimuli directions.

**Figure 3:**
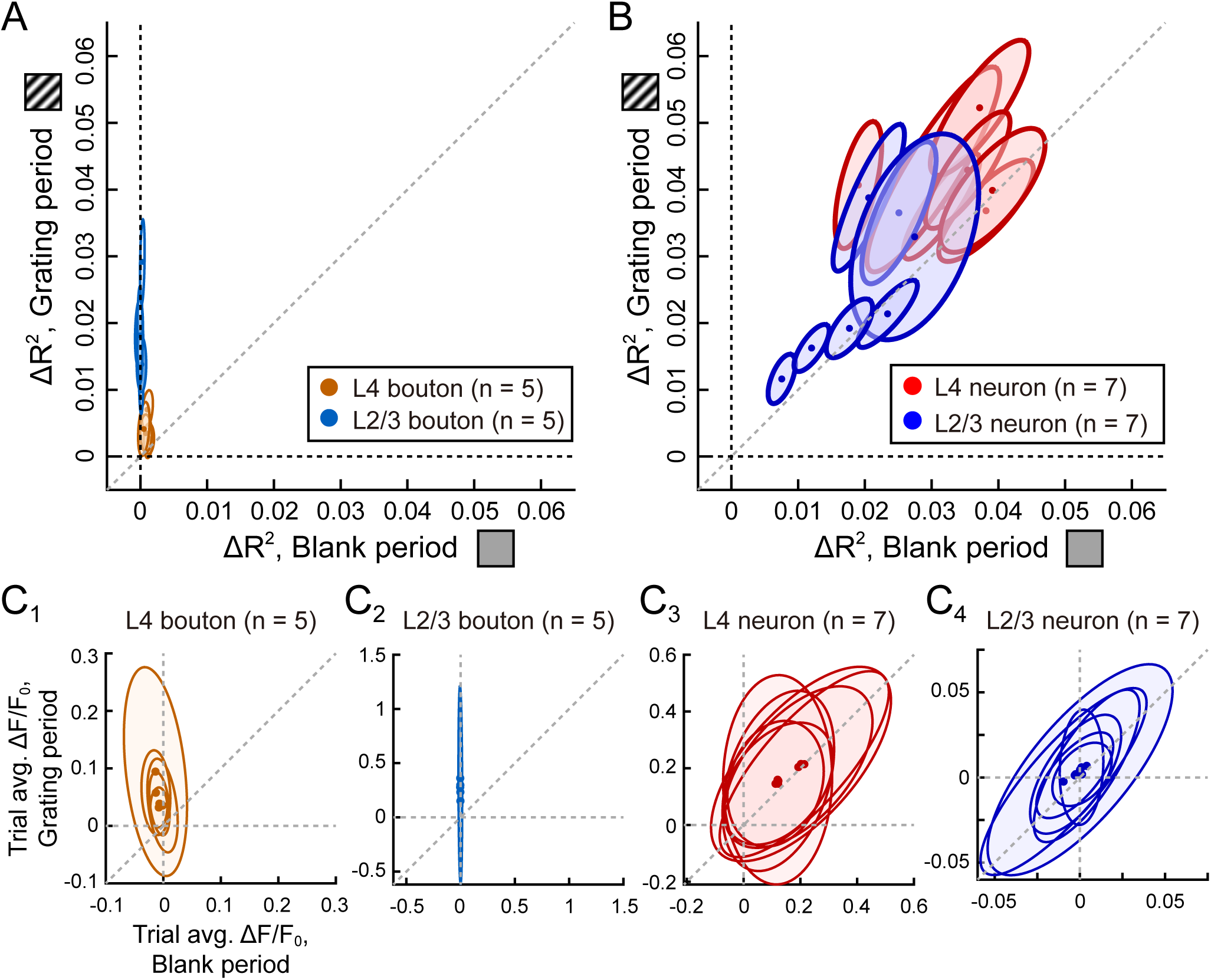
(A, B) Elevated Δ*R*^2^ correlation between facial movement and dLGN bouton (A) or cortical neuron (B) population activity during the grating stimulus period (along y-axis) compared to responses during blank stimulus period (along x-axis). Each ellipse represents the covariance of Δ*R*^2^ under the two stimulus conditions over all permutations for each dataset; the dot in the center represents the average. Different colors signify datasets in L4 or L2/3, in dLGN bouton or cortical neuron. *n* is the number of datasets used. (C) Trial average Δ*F/F*_0_ activity for dLGN bouton or cortical neuron population during the grating stimulus period (along y-axis) compared to that during blank stimulus period (along x-axis). Ellipse represents the covariance over all boutons or neurons; the dot in the center represents the average.

The activities of boutons in both L2/3 and L4 are almost uncorrelated with facial movement when blank visual stimuli are presented (Figure 3A, x axes). This is expected since the measured activity of thalamo-cortical boutons are minimal without external grating visual stimuli and therefore will not correlate with any behavioral variables (Figure 3*C*_1_, 3*C*_2_, x axes). However, surprisingly, during grating visual stimulation boutons become significantly active (Figure 3*C*_1_, 3*C*_2_, y axes) and their activity is correlated with facial movement (Figure 3A, compare x and y axes). This indicates that facial movement – which potentially reflects the behavioral state of animal – does affect the LGN feedforward visual pathway during grating stimulus periods.

For L4 and L2/3 cortical neurons, there are significant activity during both blank and grating stimuli presentations (Figure 3*C*_3_, 3*C*_4_). Consequently, there are correlations between cortical neuron activity and facial movement during both blank and grating stimuli periods in L2/3 and L4. However, we note that the correlation during grating periods is significantly higher than that during blank periods (Figure 3B, compare x and y axes). As LGN activity is uncorrelated with facial movement during blank stimulus presentation we conjecture that top-down modulation reflecting the animal’s behavioral state accounts for the facial movement correlation in V1 cortical neurons during these periods. The elevated correlation between facial movement and cortical neuron activity during grating stimulus period motivates the mechanistic analysis in the following section.

### Oculomotor signals are insufficient to account for facial movement correlation with dLGN feedforward pathway during visual stimulation

Oculomotor signals (i.e. eye movements) are a potential source of the facial movement modulation in thalamo-cortical boutons during visual stimulation. One straightforward mechanism is ‘reafference’: any behavioral correlated eye movement will create different stimulus drive, leading to behavioral correlation in feedforward inputs to V1 via thalamo-cortical boutons (Muller et al., 2023). Additionally, behaviorally driven neuromodulators could affect synapses related to eye movement, such as retinal boutons in superior colliculus (Schröder et al., 2020). Thus, it is possible that while we considered several non-eye related keypoints in our decomposition of facial movement (nose, whiskers, etc.), the correlation between dLGN bouton activity is driven primarily by eye movement.

To investigate the contribution of eye movement to the overall facial movement correlation with boutons during visual stimulation, apart from regular session permutation control (Figure 4A, Control 1), we consider an additional permutation control where the sequence of sessions for eye movement keypoints (Figure 4A, four keypoints in the blue box) are kept intact, and only all the other non-eye movement keypoint features are permuted (Figure 4A, Control 2). Thus, the session permuted control regression preserves the genuine correlation carried by eye features movement, while disrupting any correlations carried by other non-eye features. The difference, Δ*R*^2^, will contain only the unique contribution of non-eye movement features. Across bouton datasets during visual stimulation, the non-eye feature contribution is significantly above zero, yet weaker than the correlation between all face features and bouton activity (Figure 4B, compare x and y axes). Thus, oculomotor signals via eye movement explain only part of the facial motion correlation in the visually stimulated dLGN feedforward pathway. We conjecture that the residual contribution suggests alternative mechanisms independent of eye movement, such as the cortico-thalamic feedback circuit (Hasse and Briggs, 2017; Constantinople and Bruno, 2013) or other pathways.

**Figure 4:**
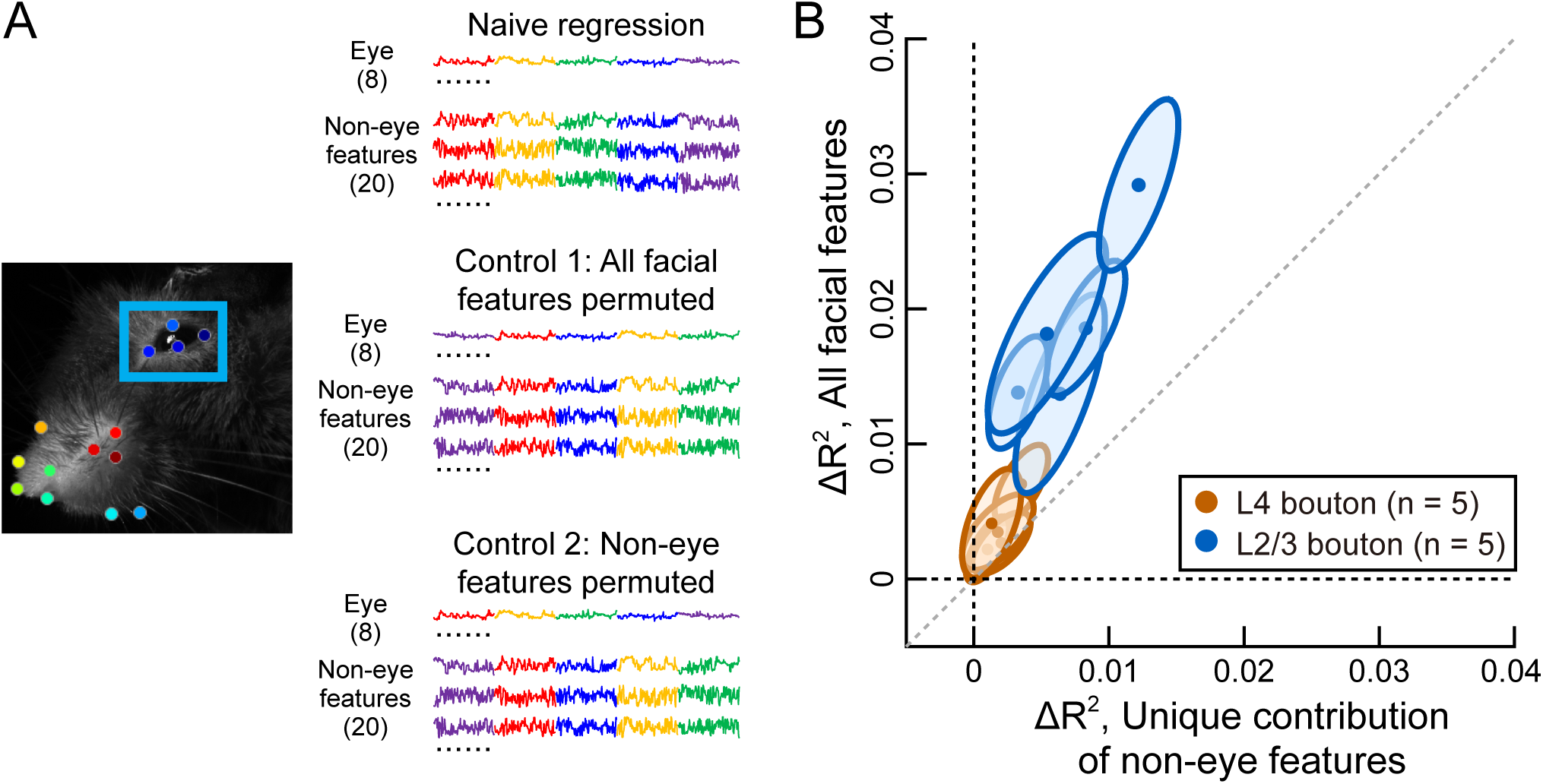
Eye movement contributes part of the correlation between facial movements and dLGN bouton activity during grating stimulus presentation. (A) During session permutations of facial movement as the control to remove nonsense correlations, we consider two groups of permutation controls: (1) permute the sequence of sessions for all facial features, so the difference Δ*R*^2^ represents the correlation of all facial features; (2) permute the sequence of sessions for only non-eye-movement features, and leave eye-movement features fixed, so that the difference also removes the genuine correlation contributed by eye movement independently, and by shared movement of eye and non-eye features, so the remainder Δ*R*^2^ represents the unique contribution of non-eye-movement. Different colors of traces indicate different sessions, as in Figure 2B. (B) For dLGN bouton activity during grating stimulus periods, the non-eye-movement contribution of facial movement continues to be correlated to dLGN bouton activity. Ellipses represent the covariance over all permutations.

### Integration of bottom-up dLGN inputs accounts for the increase of facial movement correlation with V1 neuron activity during visual stimulation

There are two potential mechanisms for elevated correlations between facial movement and cortical neuron activity during visual stimulation (Figure 3B). The first is through an increase of top-down modulatory inputs with visual stimulation, boosting the baseline correlation observed during the blank stimulus periods. The second mechanism involves a bottom-up source of facial movement signals during visual stimulation that cortical neurons combine with facial movement inputs coming from top-down pathways.

To test the first proposed mechanism we measured whether the impact of top-down modulatory input changes between blank and grating stimuli periods. To do so we considered pupil area as a proxy for arousal level which is controlled by top-down modulatory input (Raut et al., 2025). Across the datasets with valid pupil area tracking, pupil area did not differ significantly between blank and grating stimulus periods (Figure 5A). This argues against a global rise in arousal during visual stimulation as a source of increased top-down modulatory drive to cortex. We next considered a linear model between neuronal activity and the joint pupil area and facial movement dataset. We used a session permutation method to quantify the unique contributions of each factor – facial movements (Figure 5B, Näıve regression vs Control 1) or pupil area (Figure 5B, Näıve regression vs Control 2) – to the elevated correlations between neuronal activity and behavior during grating stimulus presentation. The unique contribution of pupil area to the elevation of Δ*R*^2^ was present but weak, whereas the unique contribution of facial movement was consistently stronger (Figure 5C). Together, these results suggest that the level of top-down input, as measured by arousal level, is insufficient to account for the elevated correlation between neuronal activity and facial movement during grating stimulation.

**Figure 5:**
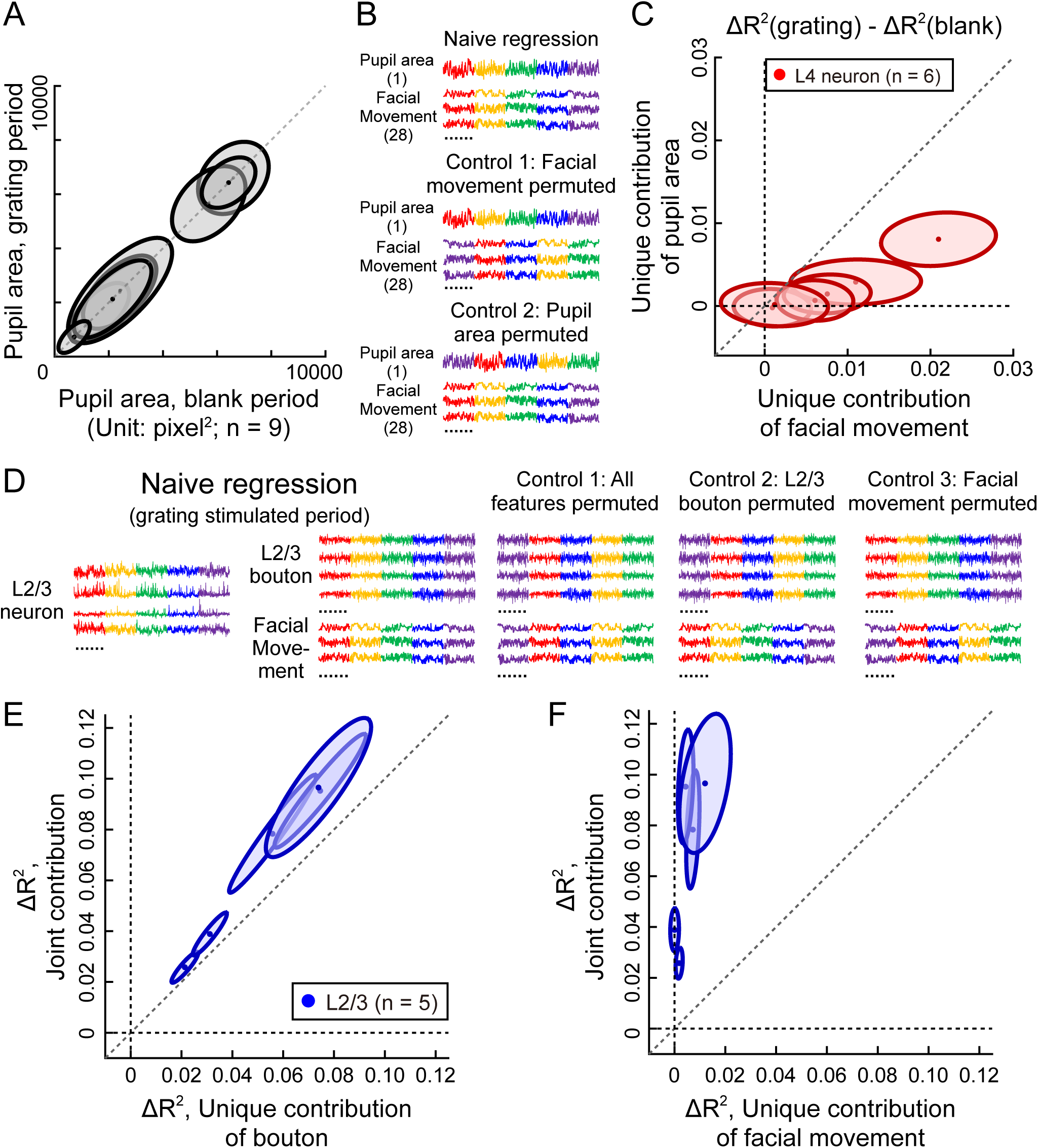
Pupil area (arousal) is insufficient to account for the elevation of facial movement correlation during grating stimulus period than blank period, while feedforward inputs explain most facial movement correlation in neurons during grating stimulus period. (A) Pupil area (proxy of the level of arousal and top-down modulatory inputs) has no systematic increase from blank period to grating stimulus period. Ellipses represent the covariance over all trials, for datasets with a tracking of pupil area. *n* = 9 represents the number of datasets with valid recording of pupil area. (B) Consider linear model between Δ*F/F*_0_ activity and a variable that concatenates pupil area and facial movement together (in different rows in the figure). Similar to control permutations in Figure 4, we consider: (1) permute the sequence of sessions for only facial movement variables and leave pupil area fixed, so the difference Δ*R*^2^ represents the unique contribution of facial movements; (2) permute the sequence of sessions for pupil area only, so Δ*R*^2^ represents the unique contribution of pupil area. Different colors of traces indicate different sessions, as in Figure 2B and 4A. (C) The elevation of Δ*R*^2^ after visual stimulation, for control groups indicating unique contribution of facial movements and pupil area. The unique contribution of pupil area to the Δ*R*^2^ elevation after visual stimulation is significantly weaker than the unique contribution of facial movement, indicating that arousal accounts for few correlation elevation after visual stimulation. Ellipses represent the covariance over all permutations. *n* = 6 represents the number of L4 neuron datasets with valid recording of pupil area. (C) For L2/3 datasets with simultaneously recording of neuron and bouton population during grating stimulus period, consider linear model between neuron Δ*F/F*_0_ activity and a variable that concatenates bouton Δ*F/F*_0_ and facial movement variables together (in different rows in the figure). Similar to control permutations in Figure 4 and 5B, we consider: (1) permute the sequence of sessions of all bouton and facial movement variable, where the difference Δ*R*^2^ represents the contribution of both to neuron activity; (2) permute the sequence of sessions of bouton activity only, where Δ*R*^2^ represents the unique contribution of bouton activity, i.e. feedforward input; (3) permute the sequence of sessions of facial movement only, where Δ*R*^2^ represents the unique contribution of facial movement but not via feedforward input. (E, F) The unique contribution of bouton activity is similar to overall contribution, while the unique contribution of facial movement is little, indicating that most facial movement correlation for L2/3 neurons during grating stimulation comes from feedforward input pathway. *n* = 5 represents the number of L2/3 neuron datasets with simultaneous recording of neuron and bouton activity.

To test the second mechanism we quantified the contribution of bottom-up inputs to the correlation between facial movement and neuronal activity during grating stimulus periods. To do so we used the subset of the datasets where simultaneous recordings of L2/3 neuron and bouton activity were acquired (*n* = 5*/*7). We considered the linear model between L2/3 neuron activity and the concatenated dataset of simultaneous bouton activity and facial movement (Figure 5D, Näıve regression vs Control 1). We can then compare the Δ*R*^2^ that are unique to either bouton activity (Figure 5D, Näıve regression vs Control 2) or facial movement (Figure 5D, Näıve regression vs Control 3), to their joint contribution. In response to grating stimuli the unique contribution from bouton activity, which represents the feedforward input, accounts for most of facial motion correlation available within the joint bouton activity and facial movement dataset (Figure 5E). By contrast, the correlation unique to facial movement but not related to feedforward input pathway is minor (Figure 5F). Thus, during grating stimulus periods most of the facial movement correlation in neuronal activity is present in (and possibly inherited from) the feedforward dLGN bouton activity. In total, this evidence suggests a convergence of a nearly persistent top-down source of behavior information (during both blank and grating stimulation), and strong behavior correlated bottom-up inputs during only grating stimulation in V1 cortical neurons.

### Decoding of visual stimuli can be separated from the encoding of facial movement in mouse V1

The analysis we have presented suggests that mouse V1 cortical neuron populations are able to encode both visual stimuli identity (direction of drifting grating for our case) and facial movement. However, facial movement does not carry any information about stimulus direction. Indeed, cross-validated linear support vector machine (SVM) decoding stimulus direction from neuron population activity during visual stimulation gives near-perfect classification accuracy, whereas simultaneous facial movement decodes at near chance (∼ 0.25, Figure 6A). Thus, we asked the question: what is the relationship between the neural representations of facial movement and stimuli direction in mouse V1? Are they largely independent from each other (as in Stringer et al., 2019), or do they overlap and potentially interfere with one another?

**Figure 6:**
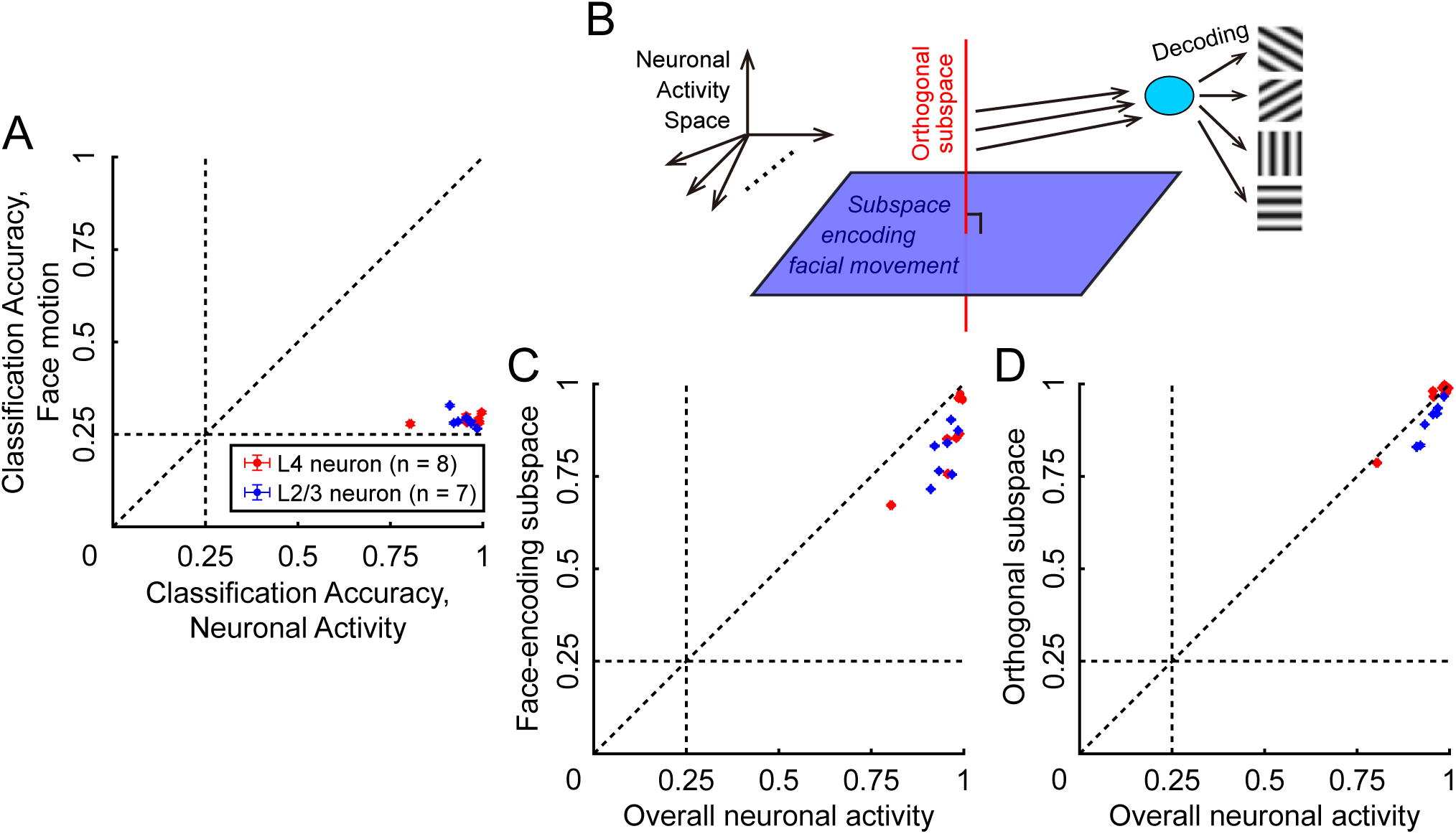
Decoding of visual stimuli information could be not interfered by the encoding of facial movement information in V1. (A) By performing the task to decode the correct visual stimuli direction from overall 4 options with linear SVM (10-fold cross-validation), the performance of neuronal activity is well, while facial movement predicts not better than by chance (1/4), indicating that face does not deliver any information of stimuli identity. Error bar represents standard deviation over 5 repeats. *n* = 7 or 8 represents the number of datasets with valid recording of grating period activity in L4 or L2/3 neuron. (B) Schematic of V1 population activity geometry. The overall neuronal activity could be separated into two components: the projection within the linear subspace encoding facial movement, and the residual orthogonal to that. (C) The activity within the facial movement subspace decodes the stimuli direction worse than the overall activity. (D) The activity orthogonal to the facial movement subspace decodes the stimuli direction equally well as the overall activity. Thus, the encoding of visual stimuli and facial movement in neuronal activity space are intermixed, since the facial movement subspace also contains information of stimuli identity; while the subspace orthogonal to that contains full information of visual stimuli for further decoding, which makes it possible to avoid the interference of facial movement modulation.

Based on the linear regression model used before, we defined the facial-movement-encoding subspace of neuronal activity and tested where stimulus information resides within the subspace. Using the same linear regression framework, we fit a mapping from facial movement features to neural activity across all directions, orthonormalized its columns to obtain the facial movement encoding subspace, and then decomposed neural activity during grating stimulus period over each direction into two components: 1) projection within this facial movement encoding subspace, and 2) its orthogonal complement (Figure 6B; see Methods). Decoding from the activity projection within the facial movement encoding subspace was well above chance but clearly weaker than that from the overall activity (Figure 6C). In contrast, the stimulus direction was decoded almost equally well from the orthogonal complement of neural activity as from the overall activity (Figure 6D).

Therefore, the encoding of facial movement and stimulus direction in mouse V1 are not fully independent from each other: stimulus information is still mostly present in the facial movement encoding subspace. However, the subspace orthogonal to the facial movement encoding subspace contains near complete information of stimuli direction for any further downstream decoding mechanism. Thus, while visual stimulus information and facial movement are intertwined in V1 population activity, stimulus identity can still be perfectly read out without interference by restricting decoding to the orthogonal subspace defined by facial movements.

## Discussion

Previous studies have shown a brain-wide modulation of cortical activity in mouse by various types of spontaneous behaviors (Stringer et al., 2019; Musall et al., 2019; Raut et al., 2025). Focusing on V1, the standard hypothesis is that modulation of cortical activity by non-visual inputs is via top-down pathways carrying information from higher level areas. However, this does not rule out that bottom-up feedforward pathways from retina through LGN also carry information about behavioral variables. To measure the contributions from bottom-up and top-down influences, we collected two-photon imaging in mouse V1 cortical neurons or thalamo-cortical boutons (i.e. feedforward inputs), as well as simultaneous tracking of facial movement. Using a ‘nonsense-free’ correlation metric, we found that bouton activity showed little or no correlation with facial movement when visual stimuli was a blank screen, but became correlated with facial movements during drifting grating stimuli presentation. This contrasted to cortical neuron activity that was correlated with facial movements during both blank and grating stimulus periods, but showed a stronger correlation when driven by grating stimuli.

We propose that during blank stimulus periods behavior information modulates V1 primarily via top-down inputs, since the dLGN pathway is weakly driven (Figure 7A). In contrast, during grating stimuli periods the dLGN pathway becomes active and is correlated with facial movement, partly through eye-motion related mechanisms (such as the motion of visual field or modulation of retinal-superior colliculus projection) and possibly through alternative routes such as potential cortico-thalamic feedback circuits (Sherman and Guillery, 2002). The convergence of dLGN bottom-up and top-down behavioral inputs in V1 during visual stimulation periods elevates the correlation between facial movements and V1 neuronal activity, compared to blank stimulus periods (Figure 7B). Finally, although facial movement carries no information about the identity of visual stimuli, the subspace of V1 population activity that encodes facial movement is not fully segregated from stimulus representations. Nevertheless, the orthogonal subspace to facial movement encoding retains full information of stimulus identity and supports decoding without interference between stimulus and behavioral variables.

**Figure 7:**
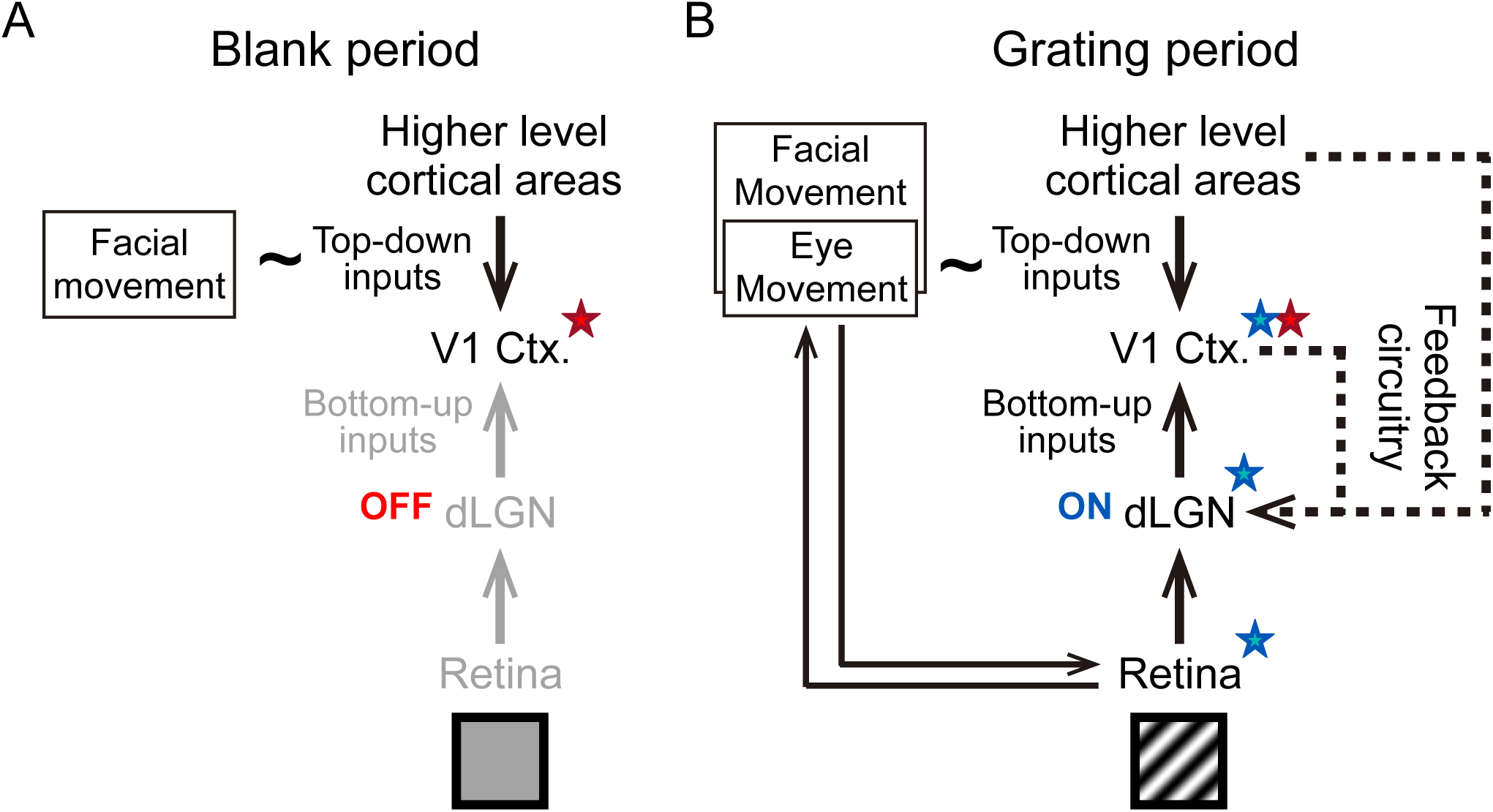
Schematic of facial movement modulation to V1 population through top-down and bottom-up pathways jointly. (A) During blank stimulus period (without grating stimulation), bottom-up (feedforward) pathway is inactivated (indicated by light gray text and arrows), and all facial movement modulation to V1 neuron population are likely due to the top-down modulatory inputs (one single red star). (B) During grating stimulus period (visually stimulated), feedforward input from dLGN (i.e. the bottom-up pathway) become correlated to facial movement (one single blue star), which is a joint work of eye movement mediated modulation and alternative, possibly feedback modulation to dLGN directly. While the top-down modulatory inputs does not become significantly stronger than that during blank period, the integration of bottom-up and top-down inputs in V1 cortex may lead to a stronger facial movement correlation (blue and red stars, as correlations from bottom-up and top-down pathways respectively).

There is a potential complication about the model framework interpretation of our dLGN bouton recordings (Figure 7). The fluorescence change in L4 and L2/3 boutons that is evoked by drafting grating stimuli is significant (Figure 3*C*_1_, 3*C*_2_, y-axes), and is also orientation tuned (Sun et al., 2016). By contrast, in response to a blank screen dLGN boutons showed minimal fluorescence change, so that a lack of movement related information in their activity is trivially expected (Figure 3*C*_1_, 3*C*_2_, x-axes). However, the relationship between calcium dynamics and bouton activity can be complicated (Ali and Kwan, 2020; Sun et al., 2016), and the bouton response to blank stimuli may be an underestimate of actual dLGN neuronal activity. Because of this the dLGN is likely not ‘off’ during blank stimulus presentation, and movement related information may indeed be transmitted. Nonetheless, extracelluar recordings of dLGN neurons in anesthetized (Durand et al., 2016; Grubb and Thompson, 2003) and awake (Durand et al., 2016) mice in response to blank or low contrast gratings visual stimuli do show very reduced firing rates, as compared to the responses to high contrast drifting gratings. This is consistent with the low activity we report for bouton activity. Further, the reduced propensity for dLGN neurons to produce bursts of spikes in response to blank compared to responses to drifting gratings (Grubb and Thompson, 2005), may also contribute to low fluoresence signals in dLGN boutons during blank stimuli periods.

One striking difference of our results from many previous reports is the weak predictive power of our linear models. Indeed, the cross-validated model never predicts more than ∼ 7% of trial-to-trial variance of neural or bouton activity from facial movement (Figure 3), far below those previously reported for either linear fits (about 10% ∼ 50% in Stringer et al., 2019) or nonlinear predictors (Syeda et al., 2024; sometimes approaching ∼ 80% in (Raut et al., 2025). Apart from differences in total trial number or animal state (mice body restrained rather than running) in our experiments, one main reason is that we explicitly estimate and remove nonsense correlations. Commonly, linear models for neuronal activity and behavioral variables use *k*-fold cross-validation to avoid overfitting, or trial shuffling as negative control to avoid chance correlations (Musall et al., 2019). However, they do not address apparent ‘nonsense’ cross-correlations that arise when two time series variables each have independent, temporally structured fluctuations. Previous studies have proposed distinct ways to estimate nonsense correlations for data with different types of temporal structures (Harris, 2020), and fortunately, our data fell into a short-term stationary regime without any drifting signals (Figure 2A), so a session permutation control was appropriate (Figure 2B, 2C). In cases where there is a drift in activity across long period recordings, identifying an adequate control is harder. One must determine whether the drift is trivial noise that should be discounted, or instead reflects a meaningful, slowly varying physiological state that should be modeled rather than subtracted (e.g., Cowley et al., 2020). Additionally, different ways to quantify correlation, the total number and time window length of trials could also affect the outcome predictive power.

More broadly, we do not claim that our linear fits provide generalizable prediction, but in practice our approach is better suited for a comparative analysis (blank vs. grating stimuli). Linear or non-linear fitting models for neurophysiological data are rarely generalizable – the fitting parameters rarely transfer across changes in stimulus sets or brain states, unless one commits to a complete mechanistic model that takes all possible conditions into account. Thus, we do not interpret the absolute magnitude of Δ*R*^2^ (in our ‘nonsense-free correlation’ metric) as predictive power. Instead, we use the linear model as a ruler to compare conditions on equal footing, such as blank period vs grating stimulus period, neuron vs bouton populations, or L4 vs L2/3; and we only interpret the differences in the Δ*R*^2^ between different conditions. Though, due to the signal-to-noise ratio of calcium imaging differs across cell types and depths (boutons vs somata, and L4 vs L2/3), we avoid drawing conclusions based on the differences between those groups. The one deliberate exception is the facial movement encoding – stimulus decoding analysis (Figure 6). Here we define a facial movement encoding subspace directly from the linear mapping, and ask how linear decoding of stimulus behaves within and orthogonal to that subspace. We argue that nonsense correlations in the linear encoding mapping does not carry information about stimulus identity, and a comparable subspace analysis is difficult to formulate in a nonlinear framework.

In our data, arousal (as indexed by pupil area) does not significantly increase from blank stimulus period to grating stimulus period, and facial movement carries no information about stimulus direction. These two observations serve as useful controls. They support our conclusion that the rise in correlation between neuronal and facial movement during grating stimulation is not simply explained by a stronger top-down drive, and that facial movement does not by itself interfere with stimulus identity decoding. These conclusions only apply to passive viewing of unrewarded, task-irrelevant stimuli, where facial movement reflects uninstructed free behavior. In task or reward contexts, arousal and facial movement may co-vary with stimulus and animal decisions. Under those conditions, both the magnitude of correlations between behavioral and neuronal activity and the overlap between neuronal subspaces encoding facial movement and stimulus identity may change. Our inferences should therefore be understood as conditional on passive, non-task visual stimulation and free behavior.

Finally, our measurements have important limitations. Direct eye gaze estimation was not available for all datasets, and high-quality eye tracking would provide a more precise measure than eye-region keypoints for isolating oculomotor contributions. Moreover, our current measurements of thalamo-cortical boutons do not link boutons to their parent LGN neurons. With identity information of boutons it would be possible to quantify redundancy information across boutons, and furthermore, to construct mechanistic theory of feedforward origin of behavioral modulation. Also, more direct measurements of top-down pathways, such as inputs into L1, would clarify how non-visual signals modulate V1. Additionally, our results suggest that a non-eye component may arrive via cortico-thalamic feedback circuitry. Testing these possibilities may be accomplished with perturbations of L6 cortico-thalamic neurons or the thalamic reticular nucleus (TRN) and asking whether bouton Δ*R*^2^ during grating stimulus period is reduced.

## STAR Methods

### Mice

All animal procedures were performed in accordance with protocols approved by the University of California, Berkeley Institutional Animal Care and Use Committee (IACUC; AUP-2020-05-13343) and complied with the National Institutes of Health guidelines for animal research. For *in vivo* imaging of the primary visual cortex (V1), adult (*>* 3 months old) mice of both sexes were used. Wild-type C57BL/6J mice (JAX 000664) were used for functional imaging of L2/3 neurons and boutons, Scnn1a-Tg3-Cre(+/+) (JAX 009613) mice were used for imaging L4 neurons, and Scnn1a-Tg3-Cre(+/-) mice were used for imaging L4 boutons. Animals were housed on a 12-hour light/dark cycles in an animal facility on UC Berkeley campus with ambient temperature controlled between 20 - 26^◦^C and humidity at 40 - 60%.

### Sample preparation: virus injection and cranial window implant

Cranial window implantation and virus injection procedures were performed following established procedures (Sun et al., 2016). All surgical procedures were conducted under 1 - 2% isoflurane in O_2_ anesthesia. A 3-mm-diameter circular craniotomy was performed over the left V1 (centered relative to midline: 2.5 mm, lambda: 1.0 mm), leaving the dura intact. Virus injection was performed using beveled glass pipettes (beveled at 45^◦^, 15-20 *µ*m opening) back-filled with mineral oil. A fitted plunger controlled by a hydraulic manipulator (Narashige, MO10) was inserted into the glass pipette, which was used to inject the viral solution into V1 or the dLGN.

To label L2/3 neurons, 30 nL of AAV9-hSyn-NES-jRGECO1a (1.8 × 10^13^ titer) was injected into WT mouse V1 300 *µ*m below the pia (9 injection sites 300 *µ*m apart). To label L4 neurons, 30 nL of AAV2/1-hSyn-FLEX-GCaMP6s (1 × 10^13^ titer) was injected into Scnn1a-Tg3-Cre mice at V1 300 - 350 *µ*m below the pia (9 injection sites 300 *µ*m apart). To label thalamic boutons in WT and Scnn1a-Tg3-Cre (+/−) mice, 30 nL of AAV2/1-hSyn-GAP43-GCaMP6s (1.2 × 10^13^ GC/ml) were injected into the dLGN (midline: 2.3 mm, lambda: 2.3 mm) at three depths (2.35, 2.55, and 2.75 mm below the pia). Following injections, the craniotomy was sealed with No. 1.5 glass coverslip using Vetbond (3M), and a titanium headbar was affixed to the skull with cyanoacrylate glue and dental acrylic.

### *In vivo* two-photon imaging

All imaging was performed at least 3 weeks after surgery on awake, head-fixed mice that were habituated to head fixation. Two two-photon fluorescence microscope systems were used. Neurons in L2/3 and L4, as well as boutons in L2/3 were imaged on a commercial two-photon microscope (Bergamo^®^ II, Thorlabs) with a Ti:Sapphire laser (Chameleon Ultra II, Coherent Inc.) tuned to 920 nm. L4 boutons were imaged on a homebuilt two-photon fluorescence system (Ji et al., 2010) with a femtosecond laser system (InSight DeepSee, Spectral Physics) tuned to 940 nm. Both setups utilized a 16×, 0.8 NA water-immersion objective (Nikon). The emitted fluorescence of GCaMP6s and jRGECO1a was detected by PMTs after transmitting through appropriate bandpass filters (Bergamo II: 525/50 nm and 607/70 nm; Custom: 510/84 nm and 617/73 nm, 630/69 nm).

Acquisition parameters were tailored to the target population. For L2/3 neurons and boutons, images (2048 × 2048 pixels; 0.3 *µ*m/pixel) were acquired at 7.6 Hz from 8 imaging planes in 4 mice at depths of 100 - 320 *µ*m below dura. For L4 neurons, images (1024 × 1024 pixels; 1 *µ*m/pixel) were acquired at 7.6 Hz from 11 imaging planes in 4 mice at imaging depths of 280 - 390 *µ*m below dura. For L4 boutons, images (300 × 300 pixels; 0.3 *µ*m/pixel) were acquired at 2.84 Hz from 6 imaging planes in 1 mouse at imaging depths of 280 - 400 *µ*m below dura. The post-objective power varied between 20 - 80 mW, depending on the imaging depth.

During all sessions, the face and pupil were recorded with an infrared-sensitive CCD camera (U-130B, Allied Vision) and infrared LED illumination. The video acquisition was synchronized to the two-photon imaging acquisition by using the same photodetector trigger and the same frame rate.

### Visual stimulation

Visual stimulus was generated using custom MATLAB^®^ scripts using Psychtoolbox (Brainard, 1997) and displayed 8 cm from the mouse’s right eye, covering 70^◦^ × 70^◦^ of visual space. Depending on the setup, a Samsung Galaxy Tablet (commercial 2PFM) or a back-projected Teflon film (McMaster-Carr) screen with a blue LED light source (450-495 nm, SugarCUBE; homebuilt 2PFM) was used. To measure the orientation tuning properties of neurons, 100% contrast circular sinusoidal drifting gratings (0.07 cycles/^◦^, 2 Hz) were presented. For variability measurements, gratings drifted in one of four directions (45, 135, 180, 270^◦^). Each direction was repeated 250 times in a pseudorandom sequence, resulting in 1000 total trials. Each stimulus lasted for 6 seconds, with a uniform gray screen displayed for the first 3 seconds and a drifting grating presented for the last 3 seconds. Stimulus and blank period onsets were marked by the appearance of a small patch of bright pixels that was detected by a photodetector (Thorlabs, PDA10A2) to trigger image acquisition and face video acquisition.

### Image processing and signal extraction

Calcium imaging data was processed by custom-written MATLAB^®^ codes. Time-lapse image sequences were registered with a rigid-motion correction algorithm (Pnevmatikakis and Giovannucci, 2017). Regions of interest (ROIs) representing individual cortical neurons were manually segmented, while ROIs representing boutons were detected and outlined for each image sequence by *Suite2p* (Pachitariu et al., 2017). For cortical neurons, the raw fluorescence signal *F* (*t*)_raw_ was extracted as the average value of the pixels within the ROI representing each neuron. To extract the neuropil signal, a circular annulus neuropil mask extending 30 *µm* from each neuron’s center (i.e. the center of mass of all pixels within the ROI) was defined, so that the neuropil signal of each neuron, *F* (*t*)_neuropil_, was the average value of pixels within the neuropil mask but outside the other ROIs identified as neurons. Then the neuropil signal was subtracted by their baseline, i.e. Δ*F* (*t*)_neuropil_ = *F* (*t*)_neuropil_ − *F*_0,neuropil_, where the baseline *F*_0,neuropil_ was defined as the most frequent mode of *F* (*t*)_neuropil_ (from the histogram count that partitions all *F* (*t*)_neuropil_ into 50 bins) for each non-overlapping time window of 25 cycles. Then, to correct the neuropil contamination in the raw fluorescence signal, Δ*F* (*t*)_neuropil_ was subtracted from raw signal *F* (*t*)_raw_, weighted by a neuropil coefficient *α*, i.e. *F* (*t*) = *F* (*t*)_raw_ − *α* · Δ*F* (*t*)_neuropil_. *α* was chosen to be 0.75 for all cells and all datasets. The baseline of corrected *F* (*t*), *F*_0_, was then defined in the same way as for *F* (*t*)_neuropil_ mentioned above. Finally, the calcium transient for each neuron was defined as Δ*F* (*t*)*/F*_0_ = (*F* (*t*) − *F*_0_)*/F*_0_. Any frames with |Δ*F* (*t*)*/F*_0_| *>* 10 were considered to be abnormal and reset as NaN. We determined whether ROIs were visually responsive by performing one-way ANOVA on the average Δ*F/F*_0_ during blank frames versus grating frames (*P <* 0.05).We included only neurons or boutons that were responsive to visual stimuli for further analysis.

For boutons, the protocol is almost the same as for neurons except the definition of baseline *F*_0,neuropil_ and *F*_0_: the most frequency mode was calculated for each individual epoch to account for moderate photobleaching of bouton fluorescence signal.

### Facial movement processing

We extracted facial keypoints from raw face videos using *Facemap*, an open-source deep-learning framework for mouse orofacial tracking (Syeda et al., 2024); we used only the pose–tracking module, which localizes predefined facial landmarks with a convolutional network. For each session, a face–centered rectangular ROI was drawn to accommodate session–to–session changes in camera pose/illumination and to exclude non–facial pixels. The base tracker was fine–tuned per session using a small set of manually annotated frames from our recordings.

We tracked *K* = 14 keypoints: four eye–orbit extrema (temporal canthus, superior, nasal canthus, inferior) and ten landmarks spanning the whisker pad, mouth, cheek, and nose. For video frame *f*, the tracker returned:

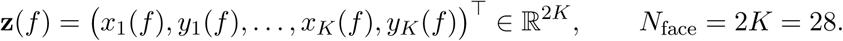

The animals’ paws were free to move, so they sometimes entered the ROI. When a paw entered the face ROI, outliers arose in two ways: (i) direct occlusion of the eye/whisker region; and (ii) out–of–distribution appearances (e.g., paw/whisker/cheek configurations not represented in the training/fine–tuning sets) that degraded predictions even without actual occlusion. To address these systematically, for each keypoint *k* and session, we flagged frame *f* as an outlier if |*x_k_*(*f*) − *µ_x,k_*| *>* 20 pixels or |*y_k_*(*f*) − *µ_y,k_*| *>* 20 pixels, where (*µ_x,k_, µ_y,k_*) are session means, and replaced flagged samples by linear interpolation between the nearest non–outlier neighbors in time (carrying the nearest valid value at segment boundaries), yielding cleaned coordinates **z̃**(*f*). For each trial *τ* containing *N*_frame_ frames, we stacked the cleaned coordinates across frames to form 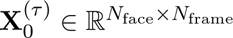, with rows ordered as (*x*_1_, *y*_1_, …, *x_K_, y_K_*].

### Linear regression analysis

For each dataset during blank or grating stimulus periods, we organized the neuronal or bouton population activity as the matrix **Y** (*N*_neuron_ or *N*_bouton_ × *N*_trial_), and facial movement activity as the matrix **X_0_** (*N*_face_ × *N*_trial_), where *N*_face_ = 28 for *x, y* locations of 14 facial keypoints (an exception is that the **X_0_** in Figure 5B contains both variable of facial movement and pupil area so there are totally 29 rows). *N*_trial_ = 1000 is consistent for all datasets – for blank stimulus periods, **Y** and **X_0_** are the averages of neuronal / bouton population activity or facial movement location, respectively, over the last several frames during the blank stimulus periods that last 400 ms, for all 1000 blank-stimulus cycles. For grating stimulus periods over the 4 different stimulus directions, **Y** and **X_0_** are the averages over 4 consecutive non-overlapping windows of the last frames during the grating stimulus period that each lasts 400 ms, for all 250 grating stimulus cycles of each direction. Note that all results during grating stimulus periods in Figures 3, 4, 5 are the average of results over 4 different directions, so that results across different stimulus directions are not intermixed.

The linear model has the form:

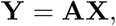

where **X** is [ones(1*, N*_trial_); **X_0_**] (the first row of 1s represents the constant offset term) and then whitened (i.e. taking the z-score), and **A** (*N*_neuron_ or *N*_bouton_ × (*N*_face_ + 1)) is the matrix of coefficient. The ordinary least-square solution of **A** that minimizes the squared sum of prediction error, i,e.

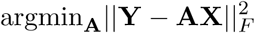

is:

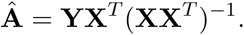

The correlation between **X** and **Y** is the *R*^2^ of the regression:

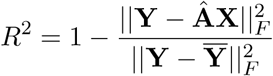

which indicates the ratio of explained trial-to-trial variance in **Y** (**Y̅** is the trial average of **Y**).

We employed 10-fold cross-validation for the linear regression and only quantify the *R*^2^ for test set. We have only 1000 trials for each dataset, which is insufficient and often leads to over-fitted training and a bad test *R*^2^. To reduce overfitting, we applied ridge regression instead which gives the following solution:

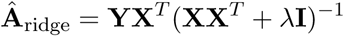

where **I** is an identity matrix and *λ* is the constant hyperparameter that determines the strength of L2 regularization to **A**. To determine *λ*, we selected the largest *λ* for which the mean test performance across (another) 10 cross-validation was within one standard error of mean of the best test performance (Semedo et al., 2019).

For the reduced rank regression in Figure 2C that determines the effective dimensionality of regression, for any given limited rank *r* for **A**, the solution is

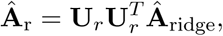

where **U***_r_* is composed of the first *r* leading principal left singular vectors of **Ŷ**_ridge_ = **Â**_ridge_**X**, where *r* can be up to 28 (number of total facial movement features). Note that all the results shown in Figures 3, 4, 5 are the asymptoted maximized Δ*R*^2^ for the reduced rank regression, after subtracting the *R*^2^ from session permutation controls.

For the extended version of linear models between Δ*F/F*_0_ activity and joint facial movement and pupil area in Figure 5, we use the matrix **X_0_** ((*N*_face_ + 1) × *N*_trial_), where the first *N*_face_ rows are facial movement and the last row is the pupil area over trials. Similarly, for the linear models between neuron activity and joint simultaneous recorded bouton activity and facial movement, the matrix **X_0_** ((*N*_face_ +*N*_bouton_)×*N*_trial_), is the concatenation of facial movement and bouton activity matrices.

### Facial movement encoding subspace analysis

To define the linear subspace of neuron population activity that encodes facial movement during visually stimulated state, we performed linear regression

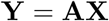

where **Y** and **X** are neuronal activity and facial movement matrices concatenated over all trials during all stimulus directions, i.e. totaling 4000 stimulated trials. **A** is computed with the same ridge regression method as described above. Consider **A** = [*b,* **A_0_**] where **A_0_** is the ‘slope’ and *b* is the constant intercept. The left singular matrix of **A_0_** (or, equivalently, the Gram-Schmidt orthogonalization of columnar vectors of **A_0_**), **Q** is the orthogonal subspace spanned by columnar vectors of **A_0_**, and it represents the subspace that linearly encodes facial movement in the visually stimulated neuronal population.

Then, the neuronal population response during each stimulus direction *k*, **Y**^(*k*)^, can be decomposed into two components: 1) the projection within the facial movement subspace:

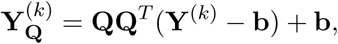

and 2) the remainder that orthogonal to the facial movement subspace:

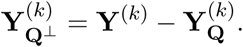

Figure 6 analyzed the linear SVM classification performance of **Y**^(*k*)^, 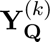, 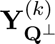, respectively.

## Data and computer code availability

All associated datasets and computer code can be publicly accessed at https://github.com/brain-math/Behavioral-correlation

## Acknowledgments

B.D and N.J. are supported by the NIH grant U19NS107613-01. B.D. is also supported by NIH grants R01NS133598 and R01EY037119 and the Simons Foundation Collaboration on the Global Brain. This research benefited from Physics Frontier Center for Living Systems funded by the National Science Foundation (PHY-2317138). Financial support was provided via the National Institute from Mathematics and Theory in Biology (Simons Foundation award MP-TMPS-00005320 and NSF award DMS-2235451). N.J. is supported by NIH grant R01NS109553 and Weill Neurohub. Y.X. is supported by a NeuroData Discovery Grant from the Kavli Foundation.

## References

Ali, F., & Kwan, A. C. (2020). Interpreting in vivo calcium signals from neuronal cell bodies, axons, and dendrites: A review. Neurophotonics, 7 (1), 011402–011402.

Aydın, C, Couto, J., Giugliano, M., Farrow, K., & Bonin, V. (2018). Locomotion modulates specific functional cell types in the mouse visual thalamus. Nature communications, 9 (1), 4882.

Brainard, D. H. (1997). The psychophysics toolbox. Spatial Vision, 10 (4), 433–436. 10.1163/156856897X00357

Busse, L. (2018). The influence of locomotion on sensory processing and its underlying neuronal circuits. e-Neuroforum, 24 (1), A41–A51.

Constantinople, C. M., & Bruno, R. M. (2013). Deep cortical layers are activated directly by thalamus. Science, 340 (6140), 1591–1594.

Cowley, B. R., Snyder, A. C., Acar, K., Williamson, R. C., Yu, B. M., & Smith, M. A. (2020). Slow drift of neural activity as a signature of impulsivity in macaque visual and prefrontal cortex. Neuron, 108 (3), 551–567.

Dolensek, N., Gehrlach, D. A., Klein, A. S., & Gogolla, N. (2020). Facial expressions of emotion states and their neuronal correlates in mice. Science, 368 (6486), 89–94.

Durand, S., Iyer, R., Mizuseki, K., De Vries, S., Mihalas, S., & Reid, R. C. (2016). A comparison of visual response properties in the lateral geniculate nucleus and primary visual cortex of awake and anesthetized mice. Journal of Neuroscience, 36 (48), 12144–12156.

Elber-Dorozko, L., & Loewenstein, Y. (2018). Striatal action-value neurons reconsidered. Elife, 7, e34248.

Erisken, S., Vaiceliunaite, A., Jurjut, O., Fiorini, M., Katzner, S., & Busse, L. (2014). Effects of locomotion extend throughout the mouse early visual system. Current Biology, 24 (24), 2899–2907.

Grubb, M. S., & Thompson, I. D. (2003). Quantitative characterization of visual response properties in the mouse dorsal lateral geniculate nucleus. Journal of neurophysiology, 90 (6), 3594–3607.

Grubb, M. S., & Thompson, I. D. (2005). Visual response properties of burst and tonic firing in the mouse dorsal lateral geniculate nucleus. Journal of neurophysiology, 93 (6), 3224–3247.

Harris, K. D. (2020). Nonsense correlations in neuroscience. biorxiv, 2020–11.

Harris, K. D., & Mrsic-Flogel, T. D. (2013). Cortical connectivity and sensory coding. Nature, 503 (7474), 51–58.

Hasse, J. M., & Briggs, F. (2017). Corticogeniculate feedback sharpens the temporal precision and spatial resolution of visual signals in the ferret. Proceedings of the National Academy of Sciences, 114 (30), E6222–E6230.

Ji, N., Milkie, D. E., & Betzig, E. (2010). Adaptive optics via pupil segmentation for high-resolution imaging in biological tissues. Nature Methods, 7 (2), 141–147. 10.1038/nmeth.1411

Keller, G. B., & Mrsic-Flogel, T. D. (2018). Predictive processing: A canonical cortical computation. Neuron, 100 (2), 424–435.

Lapanja, T., Micheli, P., Gonźalez-Guerra, A., Radomskyi, O., De Franceschi, G., Muraveva, A., Attinger, A., Roth, C. N., Tripodi, M., Boissonnet, T., et al. (2025). Pupil size modulation drives retinal activity in mice and shapes human perception. Nature Communications, 16 (1), 7334.

Liang, L., Fratzl, A., Reggiani, J. D., El Mansour, O., Chen, C., & Andermann, M. L. (2020). Retinal inputs to the thalamus are selectively gated by arousal. Current Biology, 30 (20), 3923–3934.

Marzullo, T., Rantze, E. G., Antze, R., & Gage, G. J. (2016). Stock market behavior predicted by rat neurons. Annals of Improbable Research, 12, 401.

McGinley, M. J., Vinck, M., Reimer, J., Batista-Brito, R., Zagha, E., Cadwell, C. R., Tolias, A. S., Cardin, J. A., & McCormick, D. A. (2015). Waking state: Rapid variations modulate neural and behavioral responses. Neuron, 87 (6), 1143–1161.

Meijer, G. (2021). Neurons in the mouse brain correlate with cryptocurrency price: A cautionary tale. Peer Community Journal, 1.

Muller, K. S., Matthis, J., Bonnen, K., Cormack, L. K., Huk, A. C., & Hayhoe, M. (2023). Retinal motion statistics during natural locomotion. Elife, 12, e82410.

Musall, S., Kaufman, M. T., Juavinett, A. L., Gluf, S., & Churchland, A. K. (2019). Single-trial neural dynamics are dominated by richly varied movements. Nature neuroscience, 22 (10), 1677–1686.

Nestvogel, D. B., & McCormick, D. A. (2022). Visual thalamocortical mechanisms of waking state-dependent activity and alpha oscillations. Neuron, 110 (1), 120–138.

Niell, C. M., & Stryker, M. P. (2010). Modulation of visual responses by behavioral state in mouse visual cortex. Neuron, 65 (4), 472–479.

Pachitariu, M., Stringer, C., Dipoppa, M., Schröder, S., Rossi, L. F., Dalgleish, H., Carandini, M., & Harris, K. D. (2017). Suite2p: Beyond 10,000 neurons with standard two-photon microscopy. bioRxiv. 10.1101/061507

Pnevmatikakis, E. A., & Giovannucci, A. (2017). Normcorre: An online algorithm for piecewise rigid motion correction of calcium imaging data. Journal of Neuroscience Methods, 291, 83–94. 10.1016/j.jneumeth.2017.07.031

Raut, R. V., Rosenthal, Z. P., Wang, X., Miao, H., Zhang, Z., Lee, J.-M., Raichle, M. E., Bauer, A. Q., Brunton, S. L., Brunton, B. W., & Kutz, J. N. (2025). Arousal as a universal embedding for spatiotemporal brain dynamics. BioRxiv, 2023–11.

Reimer, J., Froudarakis, E., Cadwell, C. R., Yatsenko, D., Denfield, G. H., & Tolias, A. S. (2014). Pupil fluctuations track fast switching of cortical states during quiet wakefulness. neuron, 84 (2), 355–362.

Roth, M. M., Dahmen, J. C., Muir, D. R., Imhof, F., Martini, F. J., & Hofer, S. B. (2016). Thalamic nuclei convey diverse contextual information to layer 1 of visual cortex. Nature neuroscience, 19 (2), 299–307.

Salkoff, D. B., Zagha, E., McCarthy, E., & McCormick, D. A. (2020). Movement and performance explain widespread cortical activity in a visual detection task. Cerebral Cortex, 30 (1), 421–437.

Schneider, D. M. (2020). Reflections of action in sensory cortex. Current opinion in neurobiology, 64, 53–59.

Schröder, S., Steinmetz, N. A., Krumin, M., Pachitariu, M., Rizzi, M., Lagnado, L., Harris, K. D., & Carandini, M. (2020). Arousal modulates retinal output. Neuron, 107 (3), 487–495.

Semedo, J. D., Zandvakili, A., Machens, C. K., Yu, B. M., & Kohn, A. (2019). Cortical areas interact through a communication subspace. Neuron, 102 (1), 249–259.

Sherman, S. M., & Guillery, R. (2002). The role of the thalamus in the flow of information to the cortex. Philosophical Transactions of the Royal Society of London. Series B: Biological Sciences, 357 (1428), 1695–1708.

Shimaoka, D., Harris, K. D., & Carandini, M. (2018). Effects of arousal on mouse sensory cortex depend on modality. Cell reports, 22 (12), 3160–3167.

Socha, K. Z., Couto, J., Whiteway, M. R., Hosseinjany, S., Butts, D. A., & Bonin, V. (2024). Behavioral modulations can alter the visual tuning of neurons in the mouse thalamocortical pathway. Cell Reports, 43 (12).

Spacek, M. A., Crombie, D., Bauer, Y., Born, G., Liu, X., Katzner, S., & Busse, L. (2022). Robust effects of corticothalamic feedback and behavioral state on movie responses in mouse dlgn. Elife, 11, e70469.

Stringer, C., Pachitariu, M., Steinmetz, N., Reddy, C. B., Carandini, M., & Harris, K. D. (2019). Spontaneous behaviors drive multidimensional, brainwide activity. Science, 364 (6437), eaav7893.

Sun, W., Tan, Z., Mensh, B. D., & Ji, N. (2016). Thalamus provides layer 4 of primary visual cortex with orientation-and direction-tuned inputs. Nature neuroscience, 19 (2), 308–315.

Syeda, A., Zhong, L., Tung, R., Long, W., Pachitariu, M., & Stringer, C. (2024). Facemap: A framework for modeling neural activity based on orofacial tracking. Nature neuroscience, 27 (1), 187–195.

Urai, A. E., Doiron, B., Leifer, A. M., & Churchland, A. K. (2022). Large-scale neural recordings call for new insights to link brain and behavior. Nature neuroscience, 25 (1), 11–19.

Vinck, M., Batista-Brito, R., Knoblich, U., & Cardin, J. A. (2015). Arousal and locomotion make distinct contributions to cortical activity patterns and visual encoding. Neuron, 86 (3), 740–754.

Yule, G. U. (1926). Why do we sometimes get nonsense-correlations between time-series?–a study in sam-pling and the nature of time-series. Journal of the royal statistical society, 89 (1), 1–63.

